# The role of nonspecific binding in kinase–peptide recognition revealed by molecular dynamics simulations

**DOI:** 10.64898/2026.03.30.715468

**Authors:** Yichao Wu, Yangyang Zhang, Ai Shinobu

## Abstract

Protein kinases recognize their substrates through specific short peptide motifs, and this specificity has motivated broad interest in developing peptide-based kinase inhibitors. Traditional studies have focused mainly on this specific, bound-pose interaction, whereas the nonspecific interactions that precede specific binding have received comparatively little attention. Such nonspecific interactions are transient, making them difficult to capture using either experimental techniques or conventional molecular dynamics simulations, and consequently their biological significance and functional role remain poorly understood. Here, we used two-dimensional replica exchange (gREST/REUS) to study the binding process between Abl kinase and its optimal synthetic substrate, Abltide. The simulations revealed that, in addition to the specific bound pose, the surface of Abl kinase harbors multiple nonspecific binding sites that constitute distinct encounter states, mediated primarily by electrostatic interactions. Abltide proceeds toward the binding pocket from encounter states at three of these sites. As Abltide enters the pocket, the *α*EF/*α*G/*β*11 patch plays a central role in stabilizing the resulting intermediate states, primarily through hydrophobic interactions. Within the substrate-binding site, the Abltide backbone also shifts progressively out of the register of the specific bound pose, forming backbone hydrogen bonds with the kinase that are absent from the bound pose. These findings indicate that substrate recognition by Abl kinase is encoded not only at the specific binding site but also across the surrounding kinase surface, and that peptide-based inhibitor design should consider both the specific and nonspecific binding behavior of the peptide.

**Significance:** Substrate-mimicking peptides are explored as selective inhibitors of Abl kinase, an enzyme implicated in cancer. How such peptides find their binding site is poorly understood, because the transient contacts involved are difficult to detect experimentally. Using enhanced-sampling molecular dynamics simulations, we traced how the model peptide substrate Abltide reaches its specific bound pose. Abltide can bind nonspecifically to several surface regions through mainly electrostatic interactions and then enter the binding site often via a gateway patch where hydrophobic interactions dominate. Even inside the site, part of the peptide can be shifted from its final position. Substrate recognition by Abl kinase thus extends beyond the binding site, and peptide-based inhibitor design should account for nonspecific as well as specific binding.

---

Protein kinases (PKs) are essential regulators of cellular signaling pathways, and their activity must be tightly regulated, as both overactivation and loss of activity can disrupt cellular signaling and lead to various diseases (1–3). At the molecular level, they act by binding substrate proteins and catalyzing phosphorylation at specific residues, with substrate recognition typically relying on short linear peptide motifs surrounding the phosphorylation site (4, 5). Traditionally, molecular recognition has been interpreted in terms of this specific, bound-pose interaction and its binding affinity, although binding kinetics has since received considerable attention as an additional determinant of function (6, 7). However, growing evidence suggests that the nonspecific interactions that precede specific binding also shape the binding pathway and its kinetics (8, 9). In this context, substrate discrimination depends not only on the final bound pose but also on how the substrate initially engages the kinase surface and how both partners rearrange before that pose is reached. This has direct implications for the design of peptide-based inhibitors that target the substrate-binding site of protein kinases. By mimicking natural peptide substrates, these inhibitors can achieve high specificity, reducing off-target effects and minimizing toxicity (10–12). A detailed understanding of nonspecific binding, alongside the specific bound pose, provides valuable insights into the molecular determinants of recognition and can guide the design of more effective inhibitors.

Among protein kinases, Abelson kinase (c-Abl) is a ubiquitously expressed non-receptor tyrosine kinase that localizes to both the nucleus and cytoplasm and regulates cellular processes such as cytoskeletal remodeling, cell survival, and DNA damage response (1, 13– 15). Constitutive activation of Abl kinase can lead to uncontrolled proliferation and resistance to apoptosis, thereby driving oncogenic transformation (15, 16). Targeting Abl kinase activity with small-molecule tyrosine kinase inhibitors has been clinically successful (17–19), but resistance remains a major challenge (20, 21). To overcome resistance, alternative strategies beyond ATP-competitive inhibitors have been explored (22, 23). In the 1990s, the substrate preference of Abl kinase was elucidated through peptide library screening (4), and Abltide (EAIYAAPFAKKK; Figure 1A) was identified as an optimal synthetic substrate (5), which has since served as a model for designing substrate-based inhibitors of the breakpoint cluster region–Abl (BCR-Abl) fusion protein. Abl kinase has been crystallized in complex with an Abltide analog conjugated to ATP (24), providing a stable and minimal binding interface that is well suited for mechanistic study. Structural studies (24–26) show that peptide substrates occupy the active site in a defined loop orientation (Figure 1B), aligning the phosphorylation site with the catalytic residues. In this orientation, Asp363 forms a side-chain hydrogen bond with Tyr4 of Abltide, and Phe401 engages the peptide backbone at Ala5 (Figure 1C). These hydrogen bonds define the specific bound pose of Abltide.

**Fig. 1.**
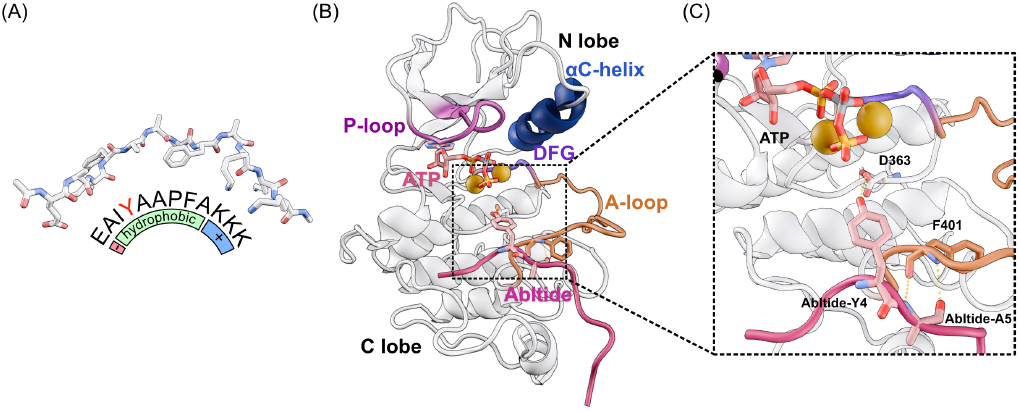
Structure of Abltide and Abl kinase. (A) The sequence and the binding structure of Abltide (EAIYAAPFAKKK), with the phosphorylatable tyrosine (Tyr4) highlighted in red. (B) The structure of the Abl-Abltide complex with ATP and Mg^2+^ ions. (C) Close-up view of the substrate-binding site. Asp363 and Phe401 of Abl kinase and Tyr4 and Ala5 of Abltide are shown in stick representation.

Only this specific bound pose, however, has been characterized, and the binding pathway and the nonspecific interactions that precede its formation remain largely unknown. Such nonspecific interactions are transient and typically too short-lived to detect with standard experimental techniques. Molecular dynamics (MD) simulations can in principle capture, at atomic resolution, these interactions and the intermediate states along the binding pathway. However, binding and unbinding events are seldom observed in conventional molecular dynamics (cMD) simulations, because the free-energy barriers separating the bound and unbound states are rarely crossed within accessible timescales (27, 28). Consequently, the biological significance and functional role of these nonspecific interactions remain poorly understood. The conformational exploration of the flexible peptide and its movement toward the substrate-binding site are slow dynamical processes that limit the ability of cMD to adequately sample the intermediate states. Capturing both processes therefore requires an enhanced-sampling strategy that can accelerate them simultaneously. Generalized replica exchange with solute tempering (gREST) (29) is a replica-exchange MD enhanced-sampling method that accelerates conformational transitions by selectively tempering interactions within targeted regions of a molecular system, enabling selective tempering of the peptide and binding-site residues. Replica-exchange umbrella sampling (REUS) (30, 31) complements gREST by exchanging umbrella potentials along a collective variable, facilitating transitions between bound, intermediate, and unbound regions. Together, gREST and REUS can be combined as a two-dimensional replica-exchange scheme (gREST/REUS) to enhance both peptide conformational flexibility and movement toward the substrate-binding site. Previous studies have applied gREST/REUS to the binding of small-molecule ligands to protein kinases (32–34). gREST has also been used to predict the pose of a peptide substrate bound to Src kinase (35).

In this study, we investigate both the specific and nonspecific binding of Abltide to Abl kinase using MD simulations. We employ the gREST/REUS method to overcome the limitations of cMD in sampling rare binding events and to capture transient nonspecific interactions. The resulting simulations provide a comprehensive free-energy landscape of the binding process, revealing multiple nonspecific encounter states and intermediate states connecting them to the specific bound pose. By resolving both the specific bound pose and the nonspecific interactions that precede its formation, our study offers an atomistic view of substrate recognition by Abl kinase. These findings provide mechanistic insights into kinase-substrate recognition and highlight the functional roles of nonspecific interactions in guiding peptide binding, offering a foundation for the rational design of peptide-based inhibitors targeting substrate-binding sites.

## Results

### Expanded exploration of the Abl-Abltide binding landscape enabled by gREST/REUS simulations

In cMD control simulations started from the bound and unbound states, Abltide sampled intermediate and encounter states, but no transitions between the bound and unbound states were observed (Figure S2). To sample these transitions, we performed gREST/REUS simulations. Effective enhanced sampling in gREST/REUS simulation requires frequent replica exchange and reversible transitions between states. We therefore assessed exchange efficiencies and bound– unbound transitions in the gREST/REUS simulations. The simulations exhibit efficient replica diffusion along both the gREST and REUS dimensions, with substantial overlap between adjacent replicas and frequent exchanges, indicating effective exploration of the parameter space (Figure S3). This effective sampling is reflected in the broad spatial distribution of the Tyr4 C*α* atom of Abltide relative to the Abl kinase at 310 K (Figure 2A). In addition, our gREST/REUS simulations successfully identified the specific bound pose (Figure 2B), indicating that the binding state is properly captured. By comparing the conformational space sampled by gREST/REUS with that from the cMD simulations, we found that gREST/REUS not only captures the transition region between the bound and unbound states but also explores areas of the landscape that were not sampled in the cMD simulations (Figure 2C). In addition, the gREST/REUS simulations exhibit continuous association and dissociation transitions. Figure 2D and Figure 2E show representative association and dissociation trajectories, respectively. These results indicate that the gREST/REUS simulations achieved wide sampling across both the bound and unbound regions.

**Fig. 2.**
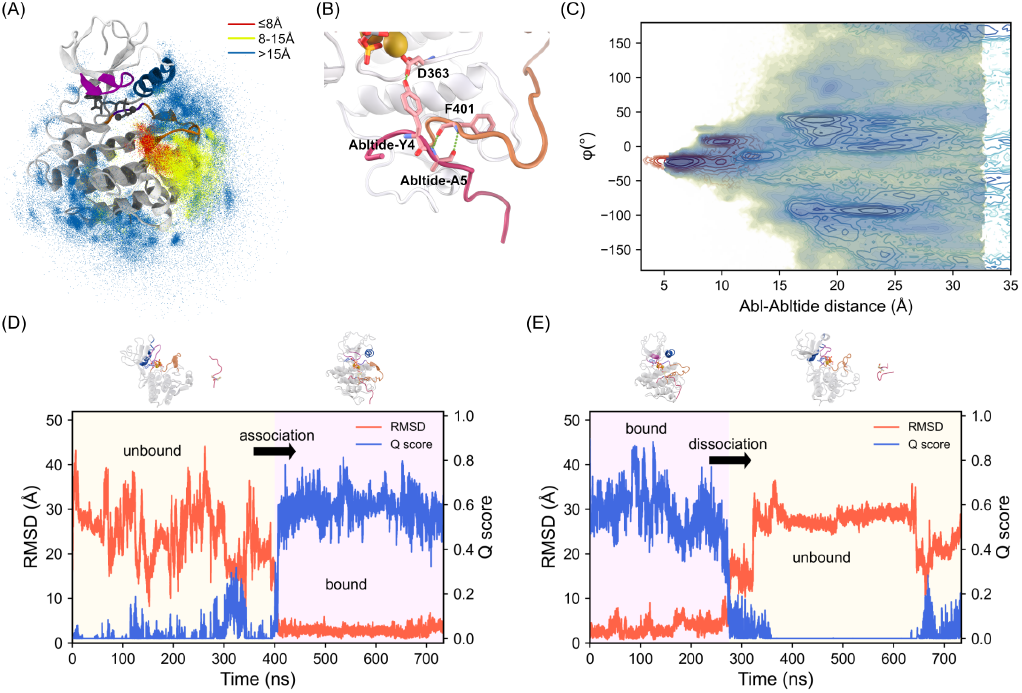
The efficiency and association/dissociation events observed in the gREST/ REUS simulations. (A) Spatial distribution of the Tyr4 C*α* atom of Abltide around Abl kinase during the gREST/REUS simulations at 310 K. Points are colored according to the Abl-Abltide distance. (B) The representative structure of the bound pose extracted from the gREST/REUS simulations. (C) Free-energy surface (FES) projected onto the Abl-Abltide distance and the spherical azimuthal angle *ϕ*. The gREST/REUS FES is overlaid with the cMD trajectory density to demonstrate the enhanced sampling achieved by gREST/REUS simulation. (D–E) Representative association (D) and dissociation (E) trajectories, shown using the root-mean-square deviation (RMSD) of Abltide (aligned to Abl kinase) and the native-contact Q score.

### Five distinct regions of encounter states identified from the free-energy surface

To characterize the binding landscape sampled by the gREST/REUS simulations, we reweighted the trajectories with the multistate Bennett acceptance ratio estimator (MBAR) (36) to construct the free-energy surface (FES). The one-dimensional free-energy profile projected onto the Abl-Abltide distance (Figure S4A) was used to define the bound (B), intermediate (I), and encounter (E) regions. The free-energy profiles obtained from the first 500 ns and the full 735 ns of each replica were similar in shape and magnitude (Figure S4B). To further characterize the spatial distribution of Abltide around Abl kinase within these regions, we performed a clustering analysis of the trajectories at the lowest solute temperature. With k-means clustering, the structures were partitioned into 25 clusters. Among these clusters, two are located in the bound region, five in the intermediate region, and 18 in the encounter regions. Intriguingly, the major barrier in the one-dimensional free-energy profile along the Abl-Abltide distance lies between the intermediate and encounter states (Figure S4A), while the encounter states are broadly distributed over the Abl kinase surface but do not lie in the binding pocket (Figure 3B). These observations suggest that the major barrier corresponds to Abltide locating the binding pocket.

**Fig. 3.**
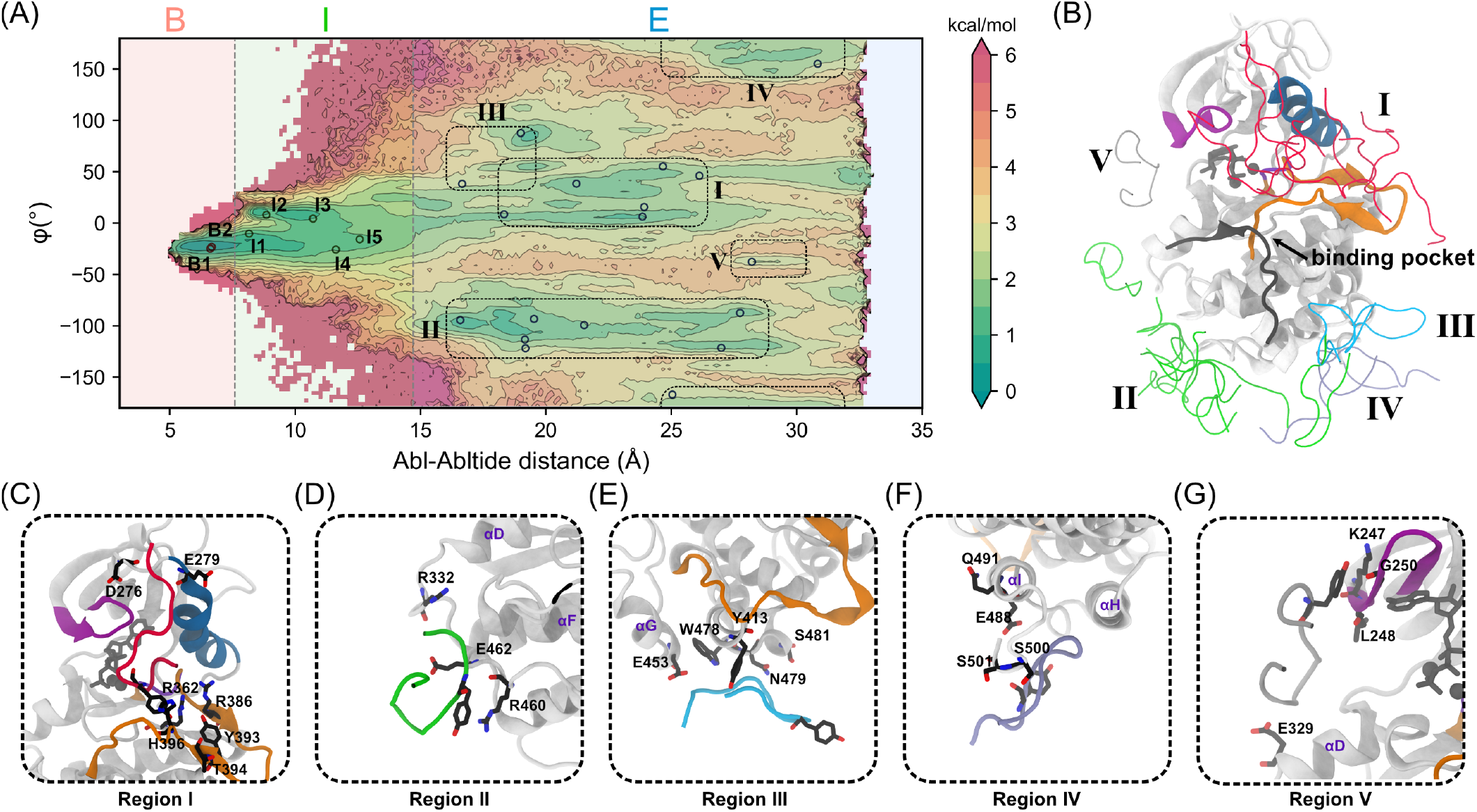
The encounter states identified from the FES. (A) FES projected onto the Abl-Abltide distance and the spherical azimuthal angle *ϕ*, divided into bound (B), intermediate (I), and encounter (E) regions. Cluster centers are overlaid on the FES, and the encounter region is further partitioned into five regions. (B) Representative cluster-center structures. Abltide in different encounter regions is shown in distinct colors, and Abltide in the specific bound pose is shown in gray. (C–G) Dominant structures of the five encounter regions. Abl kinase residues forming major contacts with Abltide and Tyr4 of Abltide, which indicates its orientation, are shown in stick representation.

We first focus on the encounter states sampled in the gREST/REUS simulations. We identified five major regions from the FES projected onto the Abl-Abltide distance and the spherical azimuthal angle *ϕ* (Figure 3A). Among these, Regions I and II correspond to the major regions of encounter states, accounting for 35.3% and 38.1% (Figure S5), respectively, and both were also observed in the cMD simulations (Figure S2). In Region I, Abltide is located near the cleft between the N- and C-lobes. In the dominant conformation of Abltide in Region I, the N-terminus of Abltide is enclosed by the activation loop (A-loop). Arg362 in the His-Arg-Asp (HRD) motif, together with Arg386, Tyr393, Thr394, and His396 in the A-loop, contributes substantially to this conformation (Figure 3C). In addition, the positively charged C-terminal segment of Abltide interacts with a negatively charged patch formed by the *β*3–*α*C loop and the N-terminal turn of the *α*C helix, in which Asp276 and Glu279 contribute most (Figure 3C, Figure S5). In Region II, as in the cMD simulations, Abltide is located on the C-lobe, near the C-terminal region of the *α*F helix (Figure 3D). In this region, Arg332, Arg460, and Glu462 contribute most to the interactions (Figure S5). In both of these major regions of encounter states, charged residues make the principal contributions to the interactions with Abltide.

In addition to these two major regions, three additional regions (III–V), which were not prominently sampled in the cMD simulations, are identified on the FES. Region III (11.2%) corresponds to a localized site on the C-lobe surface, where Tyr413, Trp478, Asn479, and Ser481 make the dominant contacts, with an additional contribution from Glu453 on the adjacent *α*G helix negative patch (Figure 3E, Figure S5). In contrast, Region IV (13.0%) is not organized around a single dominant interaction site. Abltide is instead accommodated on the C-terminal surface of the kinase domain formed by the *α*H and *α*I helices, where the contacts are distributed over many residues rather than concentrated on a few, the largest being those with Glu488 and Gln491 on the *α*I helix (Figure 3F, Figure S5). The contacts with Ser500 and Ser501 arise from the flexibility of the chain terminus rather than from a specific interaction. Region V, the least populated of the five regions (2.4%), corresponds to a site on the N-lobe, where Abltide contacts the phosphate-binding loop (P-loop) through Lys247, Leu248, and Gly250, with an additional contribution from Glu329 at the C-terminal end of the *α*D helix in the C-lobe (Figure 3G, Figure S5). These three minor regions together account for 26.6% of the encounter population, and the contacts they form are not exclusively charged. In Region III, the dominant contacts are made largely by aromatic and polar residues, whereas in Region IV, the largest contacts involve polar and charged residues in comparable measure. Considering all five regions together, the two most populated regions, which account for 73.4% of the encounter population, are dominated by charged residues, whereas aromatic and polar contacts become relatively more important in the less populated regions. Nonspecific binding of Abltide to the Abl kinase surface is thus driven predominantly by electrostatic interactions, with additional contributions from aromatic and polar residues.

### The *α***EF/***α***G/***β***11 patch stabilizes nonspecific intermediate states in the binding pocket**

After crossing the major barrier, Abltide locates the binding pocket and enters the intermediate region on the FES. In I3–I5, Abltide has not yet formed specific short-range interactions with Abl kinase (Figure S6A). Notably, the interactions in these three states are contributed by the same surface patches of Abl kinase (Figure S7). These three states are therefore treated together in the following.

The contacts formed in these states are confined to three segments of the Abl kinase surface (Figure 4A). The first comprises Lys400 and Phe401 on the *β*11 strand, which closes the A-loop and forms part of the substrate-binding site. The second comprises Leu411 and Ala412 on the *α*EF helix, together with Asn414 immediately following it. The third comprises Ser446, Tyr449, and Glu453 on the *α*G helix. Together, these residues define the *α*EF/*α*G/*β*11 patch (Figure 4C). Phe401, Leu411, Ala412, and Tyr449 stabilize the central part of Abltide, mainly through hydrophobic interactions (Figure 4B). Glu453, which lies in the *α*G helix negative patch, instead interacts with the positively charged C-terminal segment of Abltide (Figure S6F–H). Conversely, Lys400 interacts with the negatively charged Glu1 at the N-terminus of Abltide. In addition, Asn414 and Ser446 provide polar interactions. These three nonspecific intermediate states are thus stabilized primarily by hydrophobic interactions, with additional electrostatic and polar contributions.

**Fig. 4.**
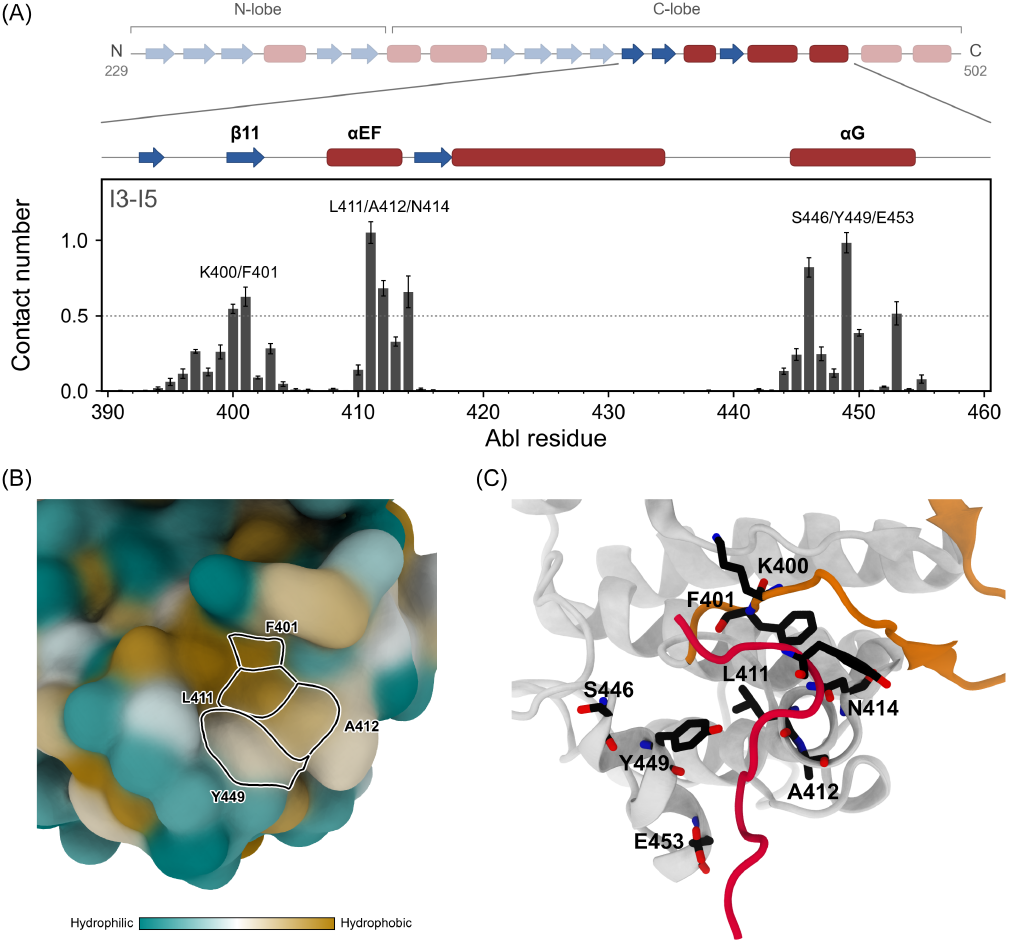
Interactions of Abl kinase with Abltide in the nonspecific intermediate states I3–I5. (A) Contact number of each Abl kinase residue with Abltide around the *α*EF/*α*G/*β*11 patch, with the secondary structure drawn above (*β*-strands as blue arrows, *α*-helices as red bars). Error bars show the standard errors estimated by block averaging (SI Appendix). (B) Hydrophobic surface representation of Abl kinase, with the surfaces of Phe401, Leu411, Ala412, and Tyr449 outlined. (C) Representative structure of I3, with Abltide in red and the A-loop in orange; the residues labeled in (A) and Tyr4 of Abltide, which indicates its orientation, are shown as sticks.

### Progressive register shift of the Abltide backbone within the sub-strate-binding site

Among the remaining clusters (Figure 3A), two lie in the bound region of the FES (B1 and B2) and two in the intermediate region (I1 and I2). B1 and B2 are therefore both regarded as bound states. In all four, Abltide lies within the substrate-binding site and forms specific short-range interactions with Abl kinase (Figure S6B–E). B1 is the state closest to the specific bound pose, as both the Asp363–Tyr4 and Phe401–Ala5 contacts are formed more frequently in B1 than in any other state, although the former is only partially formed (Figure 5B, Table S1). Phe401 is contacted mainly by Tyr4 and Ala5 of Abltide (Figure 5A).

**Fig. 5.**
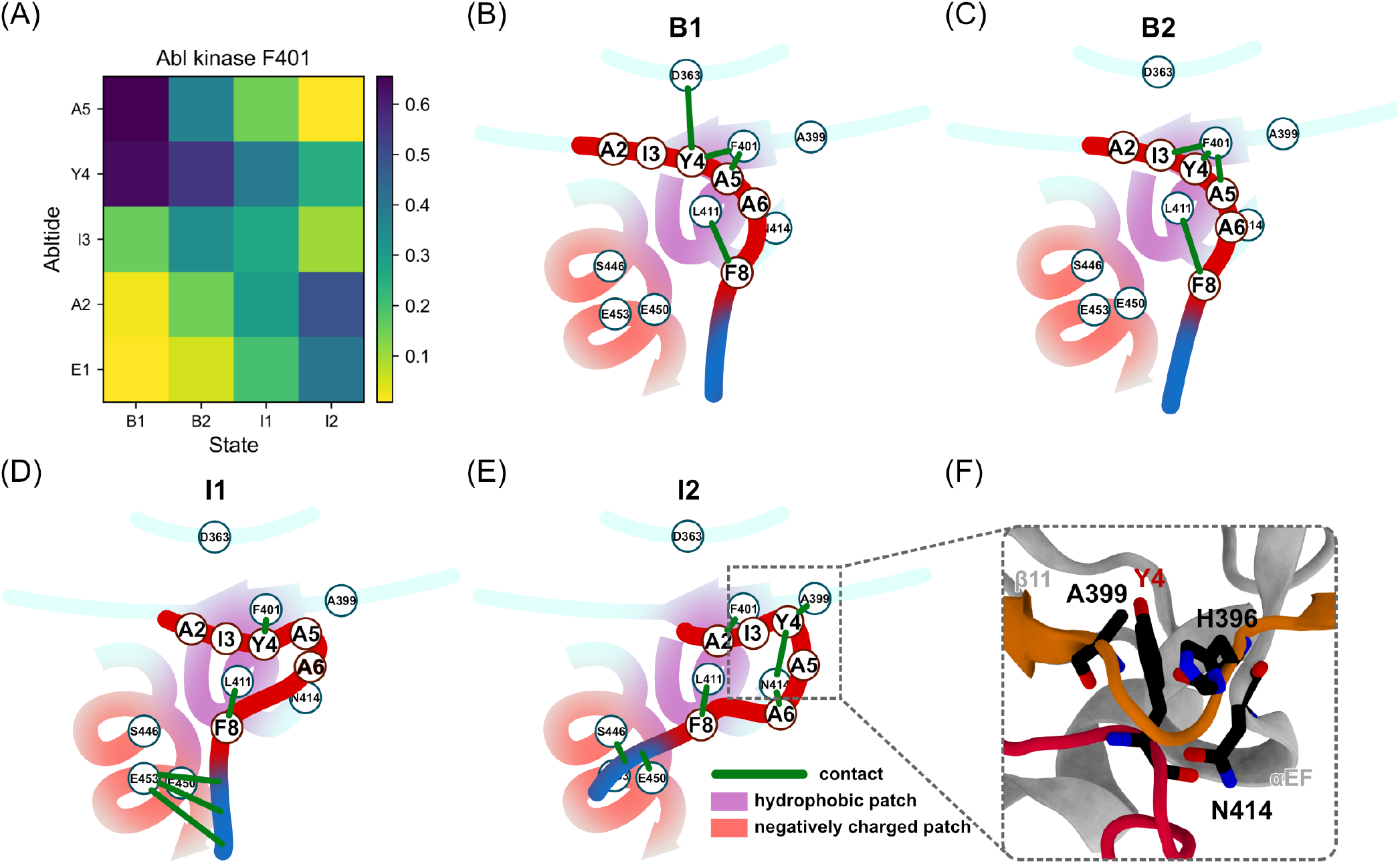
Progressive register shift of the Abltide backbone within the substrate-binding site across the B1, B2, I1, and I2 states. (A) Contact probabilities of Phe401 of Abl kinase with Glu1–Ala5 of Abltide in each state. (B–E) Schematic illustration of the interactions of Abl kinase with Abltide in B1 (B), B2 (C), I1 (D), and I2 (E). Green lines denote representative contacts between Abltide and Abl kinase. (F) Representative structure of I2 showing the region boxed in (E), with Abltide in red and the A-loop in orange; the labeled residues are shown as sticks.

In B2, the Phe401–Ala5 and Asp363–Tyr4 contacts are weaker than in B1, whereas the Phe401–Ile3 contact is stronger (Figure 5C, Table S1). The Phe401–Ile3 contact is formed not only by backbone hydrogen bonds but also by van der Waals contacts involving the backbone carbon atoms (Figure S8A, C–D). Phe401 is thus contacted by Ile3, Tyr4, and Ala5 of Abltide (Figure 5A).

In I1, neither the Asp363–Tyr4 nor the Phe401–Ala5 pair forms a specific contact, as the former is absent and the latter is weaker than in both B1 and B2 (Figure 5D, Table S1). Phe401 is instead contacted by Ala2, Ile3, and Tyr4 of Abltide with comparable probabilities (Figure 5A). In the C-terminal half of Abltide, the Leu411–Phe8 contact is enhanced, and this enhancement arises from side-chain contacts, as its backbone component remains comparable to those in B1 and B2 (Figure S8). The positively charged C-terminal segment of Abltide further contacts the *α*G helix negative patch at Glu453 (Table S1). Relative to B1 and B2, the Abltide residues contacting Phe401 shift toward the N-terminus of Abltide, which allows the C-terminal half of Abltide to form stronger contacts with Abl kinase.

In I2, the backbone of Abltide adopts a distinct register. Phe401 forms a backbone hydrogen bond with Ala2 of Abltide (Figure S8D), so that the register of the N-terminal half of Abltide is shifted by three residues relative to the specific bound pose (Figure 5E). Concomitant with this shift, the Leu411–Phe8 contact is further strengthened and, unlike in I1, is formed almost entirely by a backbone hydrogen bond between the two residues (Figure S8D). Notably, this is the only specific contact formed in all four states (Table S1), suggesting that it anchors the C-terminal half of Abltide while the register of its N-terminal half shifts. The C-terminal half of Abltide is also more extensively engaged, as Asn414 additionally contacts Ala6 and Pro7 (Table S1). The positively charged C-terminal segment of Abltide contacts Ser446 and Glu450 on the *α*G helix rather than Glu453 (Table S1). In addition, Tyr4 of Abltide forms multiple contacts, especially with Ala399 and Asn414 (Table S1). These contacts arise from the insertion of the Tyr4 side chain into the groove between the *β*11 strand and the *α*EF helix (Figure 5F). In I2, the distribution of the A-loop RMSD is also shifted to larger values relative to that in B1 (Figure S9).

Compared with the bound states B1 and B2, the Abltide residues contacting Phe401 shift toward the N-terminus of Abltide in I1 and further in I2, and the backbone hydrogen-bond partner of Phe401 changes from Ala5 in B1 and B2 to Ala2 in I2, reflecting a progressive departure of the N-terminal half of Abltide from the register of the specific bound pose (Figure 5A).

### Roles of encounter and intermediate states in the binding process

To examine how the encounter and intermediate states are connected and what roles they play in the binding process, we extracted binding paths from the continuous replica trajectories of the gREST/REUS simulations. B1 and B2 were combined into a single bound state B. Each binding path, defined as a trajectory segment that leaves the unbound state and reaches the bound state, was reduced to a loop-free path by removing excursions that return to previously visited states. Because we were interested only in the encounter region from which Abltide proceeds to an intermediate or bound state, only the last of several consecutively visited encounter regions was retained (SI Appendix). In total, 70 binding paths were identified, and the number of binding paths passing through each connection between states is shown in Figure 6.

**Fig. 6.**
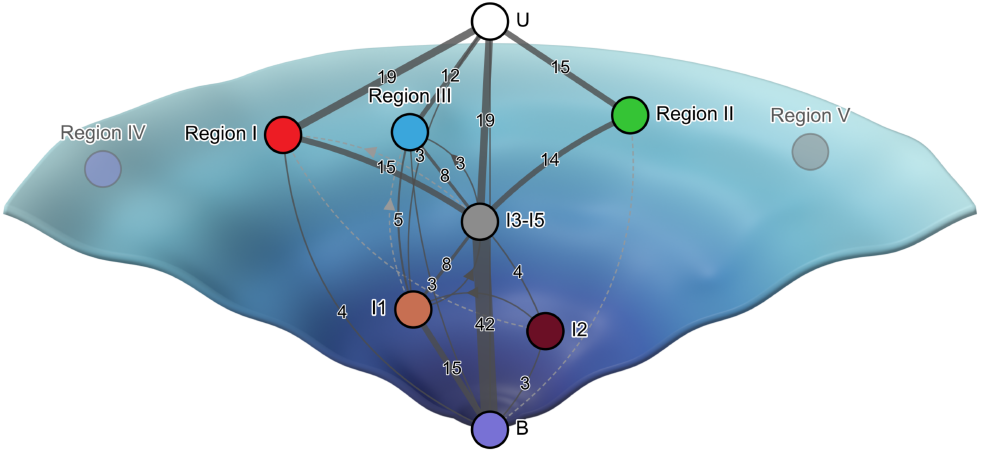
Binding pathways of Abltide to Abl kinase observed in the gREST/REUS simulations, drawn on a schematic binding funnel. The thickness of the lines represents the number of binding paths from the unbound state (U) to the bound state (B) that pass through each connection, with the corresponding number given beside each line. Connections traversed by only one binding path are shown as dashed lines. Arrows indicate connections that step back or sideways along the funnel.

The nonspecific intermediate states I3–I5 form the hub of the binding network (Figure 6). They are traversed by 58 of the 70 binding paths, whether Abltide arrives directly from the unbound state or via encounter regions. The connection traversed by the most binding paths is that from I3–I5 to the bound state. Thus, in most binding paths, Abltide associates nonspecifically with the *α*EF/*α*G/*β*11 patch before reaching the specific bound pose, indicating that this patch serves as the main gateway to the substrate-binding site.

Among the five encounter regions, Regions I, II, and III lie on the binding paths (Figure 6). From Regions I and II, Abltide proceeds mainly to I3–I5, whereas from Region III it proceeds not only to I3–I5 but also directly to I1 (Figure 6). Unlike Regions I and II, Region III involves contacts with Glu453 on the *α*G helix negative patch (Figure S5), which are also formed in I1 (Table S1). Retention of these contacts may allow Abltide to proceed directly from Region III to I1. In contrast, although Abltide occasionally visits Regions IV and V during binding, it never enters the binding pocket directly from either region.

The register-shifted states I1 and I2 also lie on the binding paths. I1 is the state within the substrate-binding site that most frequently precedes the bound state, whereas I2 does so in only a few binding paths (Figure 6). In I1, Phe401 is contacted by several residues in the N-terminal half of Abltide with comparable probabilities (Figure 5A), suggesting that this half lacks a well-defined register. Abltide in I1 may therefore adopt the specific bound pose by readjusting the register of its N-terminal half within the site. In I2, the Tyr4 side chain of Abltide inserts into the groove between the *β*11 strand and the *α*EF helix (Figure 5F). Reaching the specific bound pose from I2 may additionally involve moving this side chain out of the groove.

The binding paths analyzed here were obtained from replica trajectories along which the solute temperature and the umbrella potential change upon exchange, so that the number of binding paths passing through each connection does not represent the probability that binding proceeds through that connection at 310 K. However, because the exchanges alter only the simulation parameters while the coordinates of each replica evolve continuously, a connection that leads into an intermediate or bound state indicates that the two states are adjacent in configuration space.

## Discussion

Substrate recognition by Abl kinase involves multiple stages leading to the specific bound pose. During this process, Abltide can form encounter states through nonspecific interactions with different regions of the kinase surface. The interactions stabilizing these states vary among surface regions. Charged residues make the principal contributions in the two major encounter regions, I and II, whereas other interactions, including those involving aromatic and polar residues, also contribute substantially in Regions III–V. Analysis of the binding paths further shows that Abltide proceeds from the encounter states in Regions I–III to the intermediate states (Figure 6). Nonspecific encounter states have been proposed to accelerate association by reducing the dimensionality of the search for the binding site, from three dimensions in solution to two dimensions on the protein surface (37, 38). The encounter states in Regions I–III may therefore play this role in the binding of Abltide.

Most binding paths converge on the nonspecific intermediate states I3–I5, which form the hub of the binding network (Figure 6). In these states, Abltide associates with the *α*EF/*α*G/*β*11 patch, the main gateway to the substrate-binding site, without forming specific short-range interactions with Abl kinase. Nonspecific interactions may therefore shape the binding pathway of Abltide not only in the encounter states but also in the nonspecific intermediate states that precede the specific bound pose. As Abltide proceeds from the major encounter regions to I3–I5, its interactions with Abl kinase change from predominantly electrostatic to predominantly hydrophobic.

In the final stage of binding, Abltide forms hydrogen bonds within the substrate-binding site, and local rear-rangement of these hydrogen bonds can then yield the specific bound pose. This rearrangement can involve the backbone register of the N-terminal half of Abltide, as indicated by the register-shifted states I1 and I2 (Figure 5). Because these states precede the bound state in some binding paths (Figure 6), Abltide may enter the site in a shifted register and then reach the specific bound pose by readjusting its N-terminal half while the C-terminal half remains anchored through the interaction of Phe8 with Leu411 of Abl kinase. The shift of the A-loop RMSD distribution in I2 (Figure S9) further suggests that the binding of Abltide may be coupled to the conformation of the A-loop. The stages of Abltide binding broadly resemble those reported for peptide and protein binding, in which electrostatic interactions drive an initial nonspecific encounter with the protein surface and short-range interactions become more important as the binding partners approach the final complex (8, 9). Although charged residues dominate the major encounter regions of Abltide, the encounter regions on the binding paths also include Region III, where aromatic and polar residues make the dominant contacts (Figure 6).

Within this general progression, Abltide binds to Abl kinase through multiple heterogeneous pathways (Figure 6), a multi-state behavior that has also been identified in another protein-peptide complex and proposed to be relevant to drug design (39). An interesting question is how this heterogeneity can guide efforts to develop peptide-based inhibitors. The present results suggest two considerations for the design of peptide-based inhibitors that target the substrate-binding site of Abl kinase. First, the encounter states lie on the Abl kinase surface outside the substrate-binding site, and the residues that contribute most to the interactions with Abltide in Regions I–III, from which Abltide proceeds to the intermediate states, may serve as hot spots for designing the scaffold of a peptide-based inhibitor (Figure 6). These residues include Asp276 and Glu279, which interact with the positively charged C-terminal segment of Abltide in Region I; Arg332, Arg460, and Glu462 in Region II; and Tyr413, Trp478, Asn479, and Ser481 in Region III, together with Glu453 on the adjacent *α*G helix negative patch (Figure 3C–E, Figure S5). Second, Abltide occupies the substrate-binding site not only in the specific bound pose but also in the register-shifted states I1 and I2, in which it forms specific short-range interactions with Abl kinase other than the hydrogen bonds that define the specific bound pose (Figure 5, Table S1). In both states, Tyr4 of Abltide does not enter the catalytic pocket, so that the substrate-binding site is occupied without Abltide being positioned for phosphorylation. A peptide-based inhibitor may likewise occupy the site in a register other than that of the specific bound pose, and its design may therefore have to consider the register adopted within the site as well as the interactions formed there.

A limitation of the present study concerns the kinetic information available for the binding paths. Because the binding paths were obtained from biased replica trajectories, they cannot provide accurate kinetic information on the unbiased system. Recovering such information requires dynamical reweighting (40). One approach is to directly correct the transition probabilities estimated from these trajectories for the effect of the bias (41, 42). However, reweighting the transitions observed in the replica trajectories requires a lag time compatible with the exchange interval, a condition that is difficult to meet when the replicas are exchanged frequently. Another approach is to combine unbiased and biased simulations using multiensemble estimators (39, 43). This approach, however, relies on extensive additional unbiased simulations. Further studies are therefore needed to establish how dynamical reweighting can best be applied to gREST/REUS simulations.

Although the kinetics of binding remain unresolved, the sampled binding network shows connections among encounter states, nonspecific intermediate states, register-shifted states, and the specific bound pose. Substrate recognition by Abl kinase thus encompasses nonspecific association with the surrounding kinase surface as well as local rearrangements within the substrate-binding site.

## Materials and Methods

### System and notation

Throughout, residues of Abl kinase are numbered according to ABL1 isoform 1a, and those of Abltide from Glu1 to Lys12 (EAIYAAPFAKKK). Residue pairs between Abl kinase and Abltide are written with the Abl kinase residue first (e.g., Asp363–Tyr4). The *α*-helices and *β*-strands of Abl kinase are named according to the standard protein kinase nomenclature (26, 44) and are labeled in Figure S1. The active conformation of Abl kinase was taken from PDB entry 2G2I (24), with Pro396 mutated to histidine. The bound pose of the N-terminal eight residues of Abltide was taken from the crystallized peptide in PDB entry 2G2F (24). Phe4 of this peptide was mutated to tyrosine, and the four C-terminal residues of Abltide, which are absent from the structure, were modeled. ATP and its coordinated Mg^2+^ ions were included in the ATP-binding site. The AMBER99SB-ILDN force field (45, 46) was used for the Abl-Abltide complex, and the TIP3P model (47) was employed for water molecules. Details of the system preparation and equilibration are given in SI Appendix.

### Simulation design

All simulations were performed with GENESIS version 2.1(48), using hydrogen mass repartitioning (49, 50) to allow a time step of 3.5 fs. The cMD simulations were performed at 310 K in the NPT ensemble, with three independent 735 ns trajectories started from the bound state and three from unbound states generated by steered molecular dynamics (SMD). In the gREST/REUS simulations (29–31), Abltide and 19 binding-site residues of Abl kinase were tempered over eight solute temperatures from 310 to 481 K, while the system temperature was kept at 310 K. Umbrella potentials were applied at 36 positions along the Abl-Abltide distance, defined in SI Appendix, from 6.50 to 31.46 Å . Each of the 288 replicas was simulated for 735 ns in the NVT ensemble, for a total of approximately 212 µs. Details of the SMD, cMD, and gREST/REUS simulations are given in SI Appendix.

### Analysis

Free-energy surfaces at 310 K were obtained by MBAR reweighting (36), states were identified by k-means clustering and characterized by residue-pair contacts, and binding paths were extracted from the continuous replica trajectories, as described in SI Appendix.

### Data, Materials, and Software Availability

The input files for the SMD, cMD, and gREST/REUS simulations, together with the scripts used to analyze the simulation trajectories, will be deposited in Zenodo and made publicly available upon publication. The simulation trajectories are not deposited owing to their size and are available from the corresponding author upon reasonable request.

## ACKNOWLEDGMENTS

This work was supported by the Ministry of Education, Culture, Sports, Science and Technology (MEXT) as “Program for Promoting Research on the Supercomputer Fugaku” (Simulation- and AI-driven next-generation medicine and drug discovery based on “Fugaku”, JPMXP1020230120) and used computational resources of supercomputer Fugaku provided by the RIKEN Center for Computational Science (Project ID: hp230216, hp260023). This work was also supported by the World Premier International Research Center Initiative (WPI), MEXT, Japan. We thank Dr. Akira Imamoto for critical reading of the manuscript. During the preparation of this work the authors used Claude (Claude Opus 5.5, Anthropic) in order to improve the language and readability of the manuscript. After using this tool, the authors reviewed and edited the content as needed and take full responsibility for the content of the publication.

