## Supplementary material for "The role of nonspecific binding in kinase–peptide recognition revealed by molecular dynamics simulations": SI Appendix

**This PDF file includes:**

Supporting text
Figs. S1 to S9
Table S1
SI References

### Supporting Information Text

### Supplemental Materials and Methods

**Preparation of the Abl-Abltide complex system.** The active conformation of Abl kinase was obtained from PDB entry 2G2I (1). The bound pose of the N-terminal eight residues of Abltide on Abl kinase was taken from the crystallized peptide in PDB entry 2G2F (1). For the Abl kinase structure, missing residues were modeled using UCSF Chimera (2), and Pro396 was mutated to histidine using PyMOL (3). For the Abltide structure, PyMOL (3) was used to mutate Phe4 to tyrosine and to model the four missing C-terminal residues, which were further refined using ModLoop (4). The protonation states of histidine residues were determined using PROPKA (5, 6). The N- and C-termini of Abl kinase and Abltide were capped with neutral acetyl (ACE) and N-methylamide (NME) groups, respectively. ATP and its coordinated $Mg^{2+}$ ions were modeled from PDB entry 1ATP (7) and placed into the ATP-binding site of Abl kinase by structural alignment.

The AMBER99SB-ILDN force field (8, 9) was used for the Abl-Abltide complex, and the TIP3P model (10) was employed for water molecules. Force field parameters for ATP (11) were obtained from the AMBER Parameter Database (12). To remove any unfavorable steric clashes, the Abl-Abltide complex was first subjected to 1000 steps of energy minimization using an implicit solvent model (13, 14). After energy minimization, the structure was solvated in a rectangular box such that the minimum distance between the protein and the box boundary was 35 Å, to avoid interactions with its periodic images. A total of 59 $Na^+$ ions and 53 $Cl^-$ ions were added to neutralize the system.

We first performed 1000 steps of energy minimization while restraining the protein backbone with a force constant of 10 kcal/mol/Å$^2$ to remove steric clashes in the solvent. This was followed by 1000 steps of energy minimization without position restraints to remove steric clashes within the protein. With position restraints applied to the protein backbone, the system was then heated from 0.1 K to 310 K over 30,000 steps. Subsequently, the system was equilibrated under the NVT ensemble at 310 K for an additional 30,000 steps while backbone restraints were maintained. The restraints were then removed, and the system was equilibrated under the NPT ensemble at 310 K and 1 atm for 300,000 steps.

The following simulations were performed with GENESIS version 2.1 (15). Electrostatic interactions were calculated with the particle mesh Ewald method (16, 17). Bonds involving hydrogen atoms in the solute were constrained with RATTLE (18), and water molecules were kept rigid with SETTLE (19). Hydrogen mass repartitioning (20, 21) was applied to the solute to allow a time step of 3.5 fs with the VRES integrator (22). The temperature was controlled with the Bussi thermostat (23).

**Steered molecular dynamics simulations of the Abl-Abltide complex.** To construct the initial unbound and partially bound configurations required for the REUS component of the gREST/REUS simulations, we generated a series of starting structures using steered molecular dynamics (SMD). To prevent excessive deformation of Abltide during the pulling process and to ensure its smooth detachment from the Abl kinase, we applied geometric restraints to maintain its near-native conformation. Specifically, the C$\alpha$-C$\alpha$ distance between residues Glu1 and Lys12 of Abltide was restrained at 24.85 Å, and the angle formed by the C$\alpha$ atoms of Glu1, Ala6, and Lys12 was restrained at 95.5°. For the SMD pulling, two center-of-mass (COM) distances were defined as reaction coordinates. The first distance ranged from 6.8 to 31.3 Å with a spacing of 0.5 Å, while the second ranged from 6.7 to 31.2 Å with the same spacing. Each pulling window was simulated for 30,000 steps under the NVT ensemble at 310 K.

**Conventional MD simulations of the Abl-Abltide complex.** We performed cMD simulations starting from both the bound and unbound states. For the simulations initiated from the bound state, the initial structure was taken from the pre-equilibrated system, whereas for the simulations initiated from the unbound state, the initial structures were obtained from the final configurations of the SMD simulations. The production simulations were performed in the NPT ensemble at 1 atm and 310 K, with the pressure controlled by the Bussi barostat (24). For each system, three independent trajectories were generated, with each trajectory simulated for 735 ns. Coordinates were saved every 3000 steps (10.5 ps).

**gREST/REUS simulation of the Abl-Abltide complex.** We performed gREST/REUS simulations, which combine replica exchange along two dimensions, namely solute tempering (gREST) (25) and umbrella sampling along the Abl-Abltide distance (REUS) (26, 27). For the gREST dimension, the Coulomb, Lennard-Jones, and dihedral interaction terms of Abltide and 19 binding-site residues of the Abl kinase were selected as the solute region. The 19 binding-site residues were Asp325, Asp363, Arg367, Ala399, Lys400, Phe401, Pro402, Ile403, Trp405, Leu411, Ser438, Asp444, Leu445, Ser446, Gln447, Tyr449, Glu450, Glu453, and Lys454. Eight replicas were employed, with temperatures set to 310, 328, 349, 371, 396, 422, 450, and 481 K, respectively. For the REUS dimension, the initial structures were obtained from SMD simulations. The umbrella potentials were applied to the Abl-Abltide distance, defined as the distance between the center of mass of the C$\alpha$ atoms of Abltide and that of the substrate-binding site of Abl kinase. The substrate-binding site was represented by the C$\alpha$ atoms of residues Asp363, Arg367, Ala399, Lys400, Phe401, Pro402, Ile403, Trp405, Leu411, Ser438, Leu445, Ser446, Gln447, Tyr449, Glu450, Glu453, and Lys454, which interact with Abltide in the specific bound pose. A total of 36 replicas were employed, with umbrella centers placed at intervals of 0.65 Å to 0.75 Å along the Abl-Abltide distance from 6.50 Å to 31.46 Å, and a spring constant of 2 kcal/mol/Å$^2$. Therefore, the gREST/REUS simulations consisted of a total of 288 replicas. Exchanges were performed alternately along the gREST dimension and the REUS dimension, with exchanges attempted every 600 steps. The gREST/REUS simulations were performed in the NVT ensemble, with the thermostat temperature set to 310 K. Each replica was simulated for 735 ns, and coordinates were saved every 3000 steps (10.5 ps). The total simulation time over the 288 replicas was approximately 212 µs.

**Constructing the free-energy profile.** To characterize the binding process of Abltide to Abl kinase, we constructed a two-dimensional free-energy surface using representative reaction coordinates. One of the reaction coordinates was the Abl-Abltide distance used in REUS.

To characterize the spatial distribution of Abltide on the surface of Abl kinase, we defined another reaction coordinate. The reaction coordinate was defined as the azimuthal angle ($\phi$) and polar angle ($\theta$) of the C$\alpha$ atom of the Tyr4 residue of Abltide in a spherical coordinate system constructed on Abl kinase. Specifically, the spherical coordinate system was constructed on Abl kinase by defining the C$\alpha$ atom of Trp423 as the origin. The vector from the C$\alpha$ atom of Trp423 to that of Lys419 was defined as the y-axis. The x-axis was defined as the cross product between this vector and the vector from the C$\alpha$ atom of Trp423 to that of Phe425. The z-axis was then determined to complete a right-handed coordinate system.

The trajectories obtained from cMD and gREST/REUS simulations were projected onto these reaction coordinates. The trajectories from the gREST/REUS simulations were reweighted using the multistate Bennett acceptance ratio estimator (MBAR) (28), as implemented in PyMBAR.

To examine how the free-energy profile depends on the simulation time, the MBAR reweighting was repeated using progressively longer portions of the gREST/REUS trajectories, taking the first 10, 50, 100, 250, 500, and 735 ns of every replica, and the resulting one-dimensional profiles along the Abl-Abltide distance were compared (Figure S4B).

**Obtaining structural ensembles from clustering.** K-means clustering was performed to characterize the conformational states of Abltide during the binding process, including bound, intermediate, and encounter states. Clustering was carried out using the scikit-learn package (29). The trajectories at the lowest solute temperature (310 K) in gREST were used for the analysis. Prior to clustering, all structures were aligned using the backbone heavy atoms of Abl kinase. The C$\alpha$ atoms of both Abl kinase and Abltide were then used as input features for clustering. The number of clusters was set to 25, and the centroid structure of each cluster was used as its representative structure. The resulting clusters were classified into bound, intermediate, and encounter states based on their positions along the Abl-Abltide distance on the free-energy surface (Figure S4A). For the encounter states, the clusters were further grouped into five regions according to the local minima on the two-dimensional free-energy surface.

**Contact analysis.** Contacts were evaluated for the structures at the lowest solute temperature in the gREST/REUS simulations, each assigned to a state by the clustering described above. Two residues were considered to be in contact when the closest distance between their heavy atoms was within 3.5 Å, and the contact probability of a residue pair in a given state was calculated as the MBAR-weighted fraction of structures assigned to that state in which the contact was formed (Figure S6B–H). For the states in the bound and intermediate regions, a residue pair was considered to form a specific contact when its contact probability exceeded 0.3 (Figure S6A, Table S1). The intermediate states I3–I5, in which no residue pair formed a specific contact, were classified as nonspecific intermediate states. The contacts formed in nonspecific binding (in the encounter regions and the nonspecific intermediate states) were characterized by the contact number of each Abl kinase residue, defined as the number of Abltide residues in contact with that residue, averaged with MBAR weights over the structures assigned to a given region or state. For nonspecific binding, Abl kinase residues with a contact number above 0.5 were labeled. The standard errors were estimated by block averaging, dividing the 735 ns trajectories of the 36 umbrella windows at the lowest solute temperature into five consecutive 147 ns blocks, recalculating each quantity in each block with the MBAR weights obtained from the full trajectories, and taking the standard deviation over the blocks divided by the square root of the number of blocks. To decompose the contacts into side-chain and backbone components (Figure S8), the same distance criterion was applied separately to the side-chain heavy atoms, the backbone heavy atoms, and the backbone N and O atoms of both residues. As a simple criterion, a contact evaluated over the backbone N and O atoms was regarded as a backbone hydrogen bond, without distinguishing hydrogen-bond donors from acceptors or applying an angular criterion.

**Analysis of transition pathways.** Binding pathways were analyzed along the 288 continuous replica trajectories of the gREST/REUS simulations. Each of these trajectories follows the coordinates of a single replica while its simulation parameters are exchanged, and frames were saved every 10.5 ps. Each frame was labeled with one of the 25 clusters described in *Obtaining structural ensembles from clustering* above, and the clusters were combined into 10 states. These states were the unbound state (U), Regions I–V of the encounter states, a single state comprising the nonspecific intermediate states I3–I5, the intermediate states I1 and I2, and a single bound state (B) comprising B1 and B2, both of which lie in the bound region of the free-energy surface.

To prevent fluctuations across cluster boundaries from appearing as transitions, the trajectories were discretized using core sets (30). A frame was assigned to the core of U when the minimum heavy-atom distance between Abltide and Abl kinase exceeded 10 Å. Otherwise, a frame labeled with cluster $k$ was assigned to the core of the state containing cluster $k$ when two conditions were met. First, its distance to the center of cluster $k$ was at most 0.8 times its distance to the nearest other cluster center, where the distance to a cluster center was evaluated as the root-mean-square deviation (RMSD) of the 286 C$\alpha$ atoms of Abl kinase and Abltide from the MBAR-weighted average structure of that cluster. Second, the minimum heavy-atom distance between Abltide and Abl kinase was within 3.5 Å. Because B comprises two clusters, the first condition was applied to frames labeled with B1 or B2 by comparing the distance to the nearer of these two cluster centers with the distance to the nearest of the other 23 cluster centers. These frames were assigned to the core of B only when Abltide also adopted the pose of B1, that is, when the RMSD of the C$\alpha$ atoms of Ile3–Pro7 of Abltide from the corresponding atoms of the MBAR-weighted average structure of B1 was within 3.5 Å. This RMSD was evaluated after superposition on Abl kinase only, so that it reflects the position of Abltide on Abl kinase. Frames outside any core were assigned to the most recently visited core.

A binding event was defined as a trajectory segment that leaves the core of U and reaches the core of B without returning to U, and the sequence of states visited within this segment was taken as its binding path. Each binding path was then reduced to a loop-free path in three steps.

1. When Abltide visited several encounter regions in succession, only the last of them was retained, because the encounter region of interest was the one from which Abltide proceeds to the next stage of binding.

2. Loops were removed in chronological order; that is, whenever a state was revisited, the states visited since its previous visit were discarded.

3. The same rule was applied to encounter regions that became consecutive after loop removal.

Because the earlier encounter regions were discarded, a connection that leads into an encounter region may bypass other encounter regions visited before it. In total, 70 binding events were identified, and Figure 6 of the main text shows the number of loop-free binding paths passing through each connection between states. Because the trajectories were generated under replica exchange, these numbers characterize the connectivity among the states rather than the transition rates between them.

The scripts used to extract and analyze the binding paths were written with the assistance of Claude Code (Claude Opus 5.5, Anthropic). These scripts were carefully reviewed by the authors, who take full responsibility for the analysis.

### Supplemental Results

**Encounter and intermediate states of Abltide observed in cMD simulations.** We first performed cMD simulations starting from both the bound and unbound states. We constructed a spherical coordinate system centered on Abl kinase to analyze the binding positions of Abltide. The construction of this coordinate system is described in *Constructing the free-energy profile* above. None of the six trajectories crossed between the bound and unbound states, although Abltide deviated substantially from its starting configuration within each region. In the simulations initiated from the bound state, two intermediate states (cI1 and cI2) were identified (Figure S2). In both the cI1 and cI2 states, the Asp363–Tyr4 and Phe401–Ala5 contacts, which existed in the bound experimental structure, are disrupted (Figure S2C–D). In the cI2 state, new backbone hydrogen bonds form between Phe401 of Abl kinase and Ala2 of Abltide. This suggests that Abltide binding may involve transient interactions that are absent in the fully bound pose. For the simulations initiated from the unbound state, two trajectories were eventually captured in distinct encounter states (cE1 and cE2, Figure S2). In the cE1 state, Abltide approaches the C-terminal region of the $\alpha$F helix (Figure S2E), while in the cE2 state, it engages the cleft region (Figure S2F). These results suggest that Abltide binding involves multiple intermediate and encounter states. However, a complete transition between the bound and unbound states was not observed, indicating that cMD alone is insufficient to fully resolve this transition and motivating the use of enhanced-sampling simulations.

 

157 ## Supplemental Figures

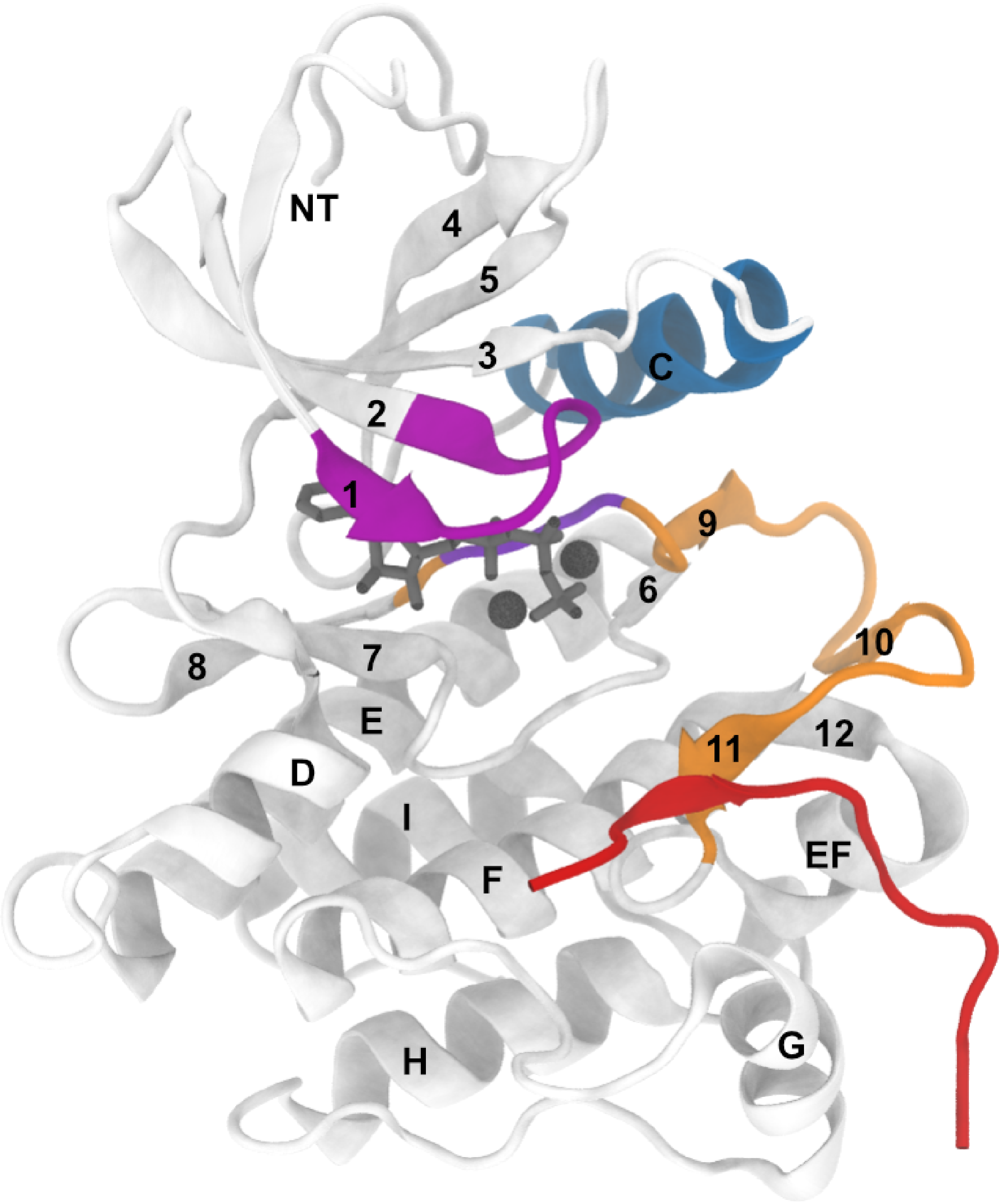


**Fig. S1.** Cartoon representation of the Abl kinase–Abltide complex. $\beta$-strands are numbered and $\alpha$-helices are labeled with letters.

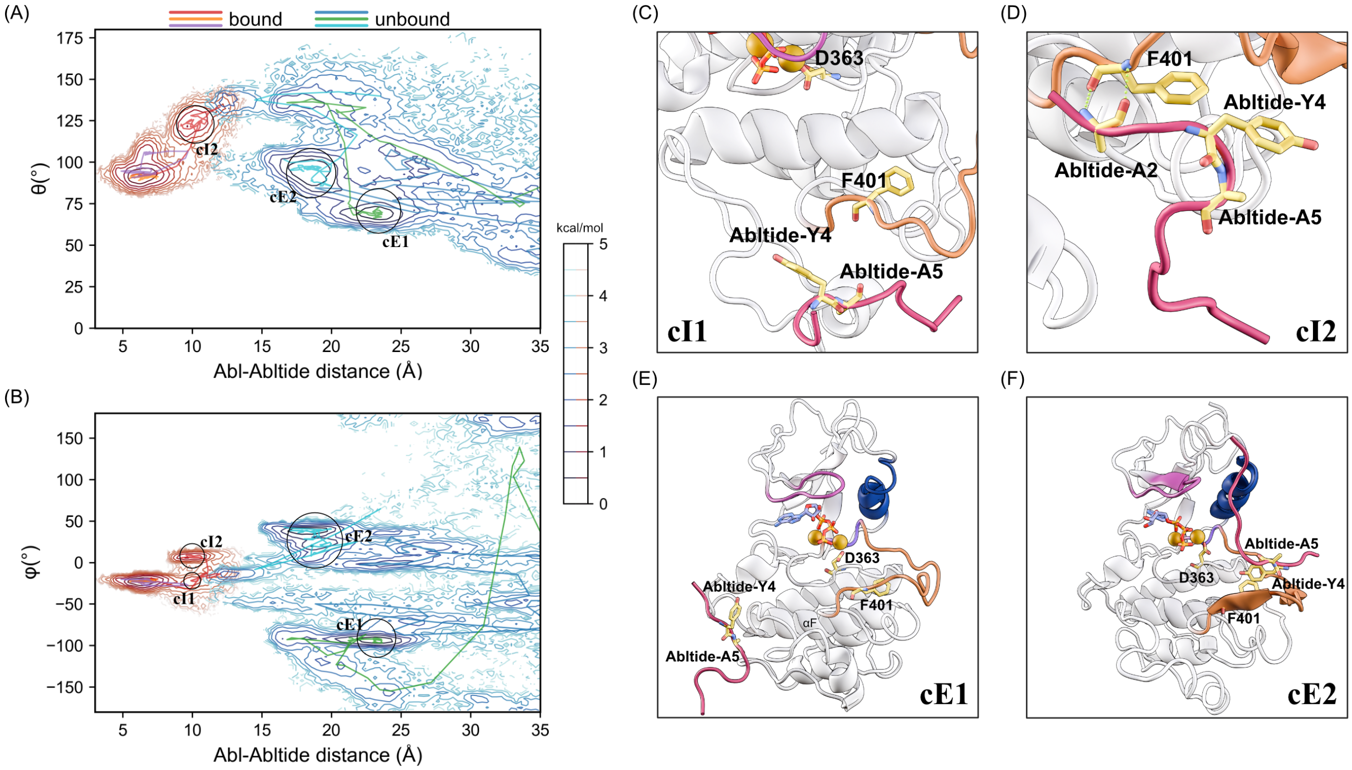


**Fig. S2.** Results of the cMD simulations of the Abl-Abltide complex. (A–B) The cMD trajectories projected onto the Abl-Abltide distance and the spherical polar angle $\theta$ (A) and azimuthal angle $\phi$ (B). Trajectories initiated from the bound state are shown in red, and those initiated from the unbound state are shown in green. (C–D) Cartoon representations of representative structures of the intermediate states cI1 (C) and cI2 (D). (E–F) Cartoon representations of representative structures of the encounter states cE1 (E) and cE2 (F).

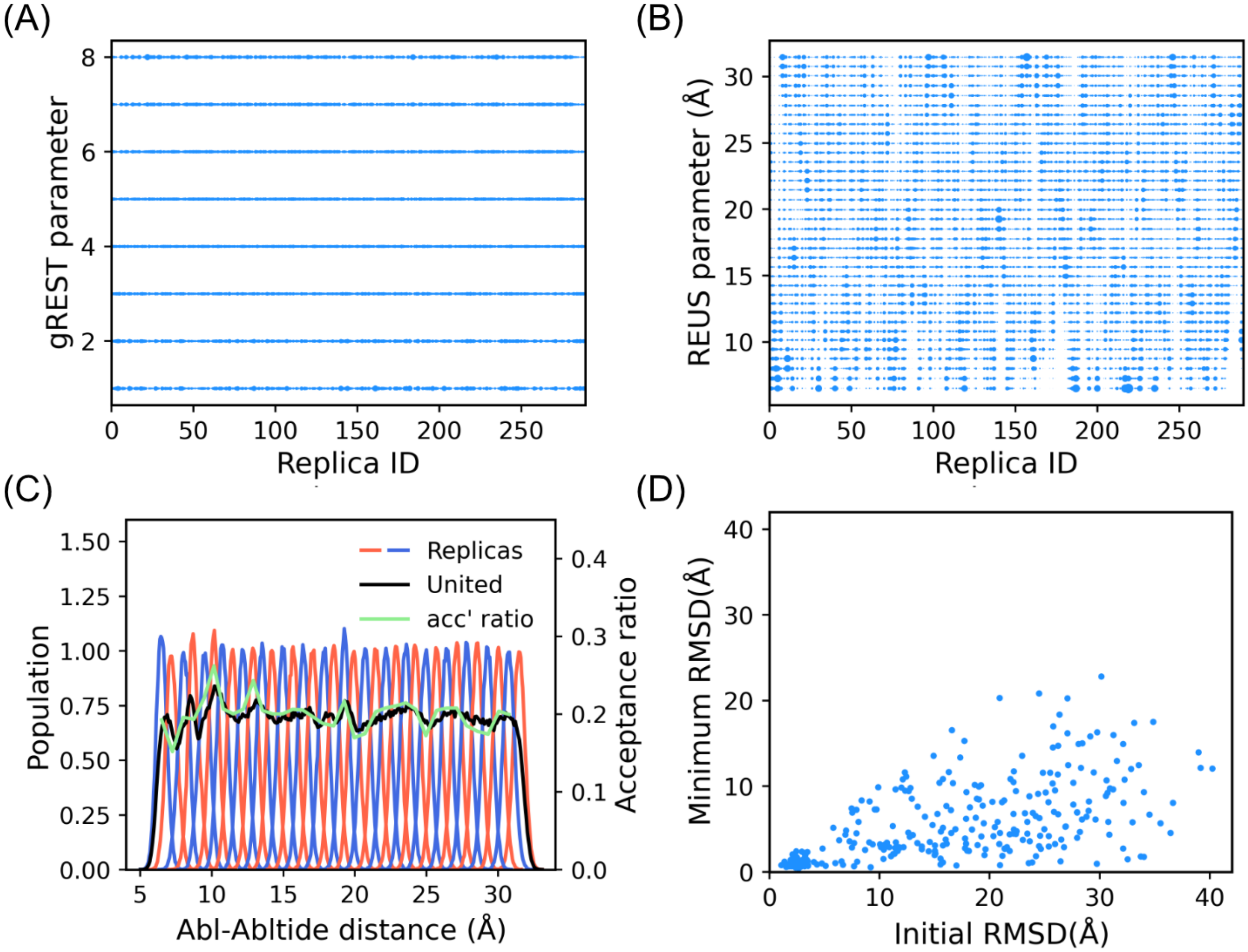


**Fig. S3.** Sampling efficiency of the gREST/REUS simulations. (A and B) Relative population of each replica at each gREST parameter (A) and at each REUS parameter (B), with the sphere size proportional to the population. (C) Distributions of the Abl-Abltide distance sampled with each REUS parameter at the lowest solute temperature (red and blue lines), together with the distribution over all REUS parameters (black line, “United”) and the acceptance ratios of exchanges between adjacent REUS parameters (green line). (D) Minimum RMSD of each replica during the simulation, plotted against its initial RMSD.

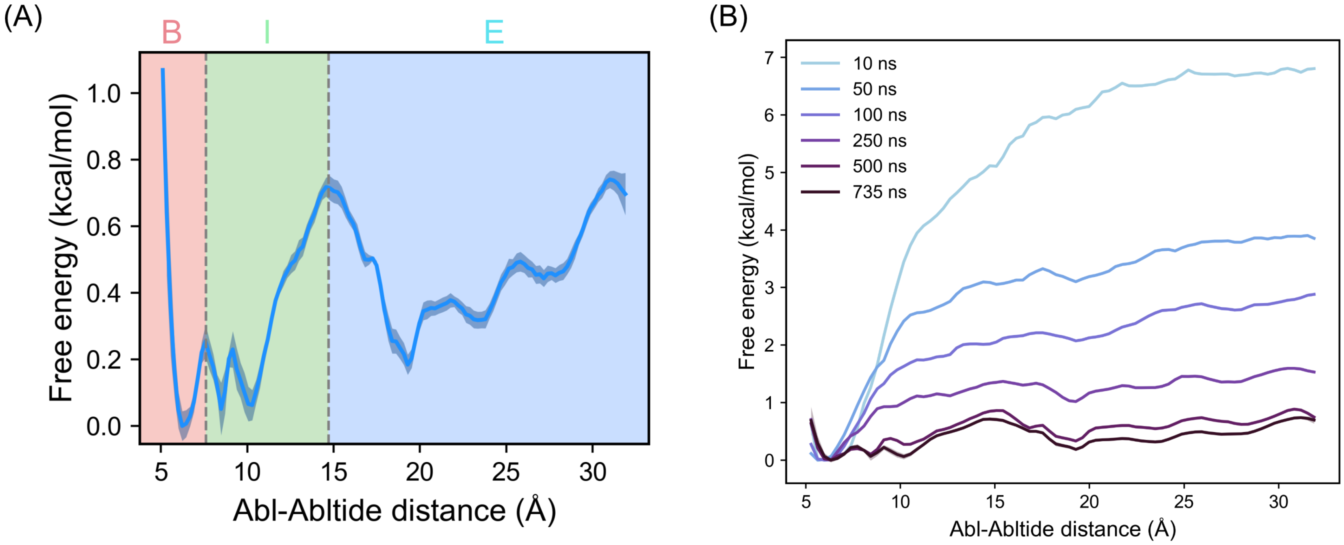


**Fig. S4.** The free-energy profile along the Abl-Abltide distance. (A) The profile obtained from the full gREST/REUS trajectories. The bound (B), intermediate (I), and encounter (E) regions are identified. The shaded band shows the standard error estimated by block averaging over five consecutive 147 ns blocks, with the MBAR weights obtained from the full trajectories. (B) Dependence of the profile on simulation time. Profiles were reweighted from the first 10, 50, 100, 250, 500, and 735 ns of every replica. The profiles obtained from 500 ns and 735 ns are nearly superimposed.

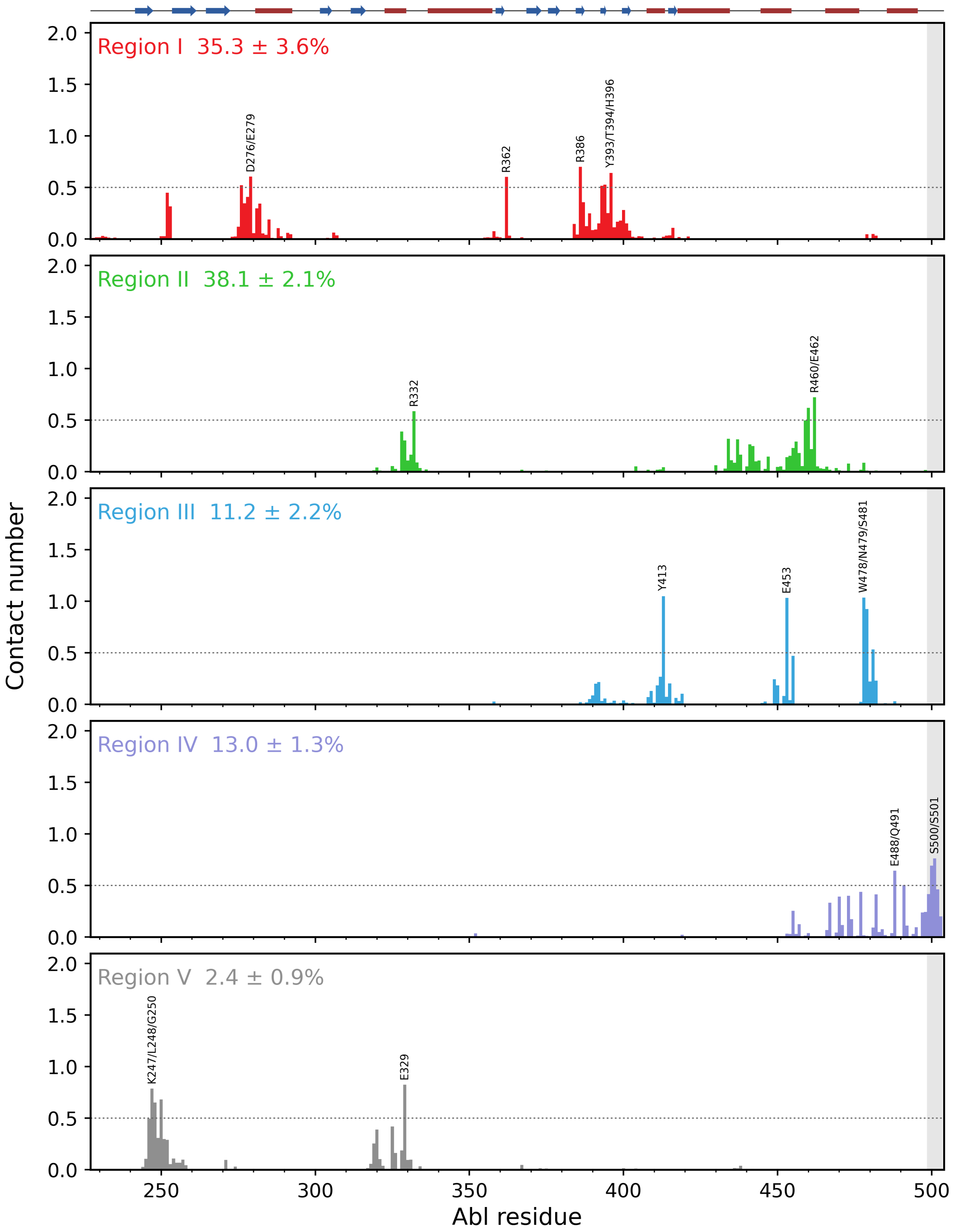


**Fig. S5.** Contact number of each Abl kinase residue with Abltide in each of the five regions of encounter states. Residues with a contact number above 0.5 are labeled. The shaded region marks residues 499–503, where the contacts seen in Region IV reflect the flexibility of the chain terminus rather than a specific interaction. The percentage in each panel gives the MBAR-weighted population of the region relative to all five regions, with its standard error.

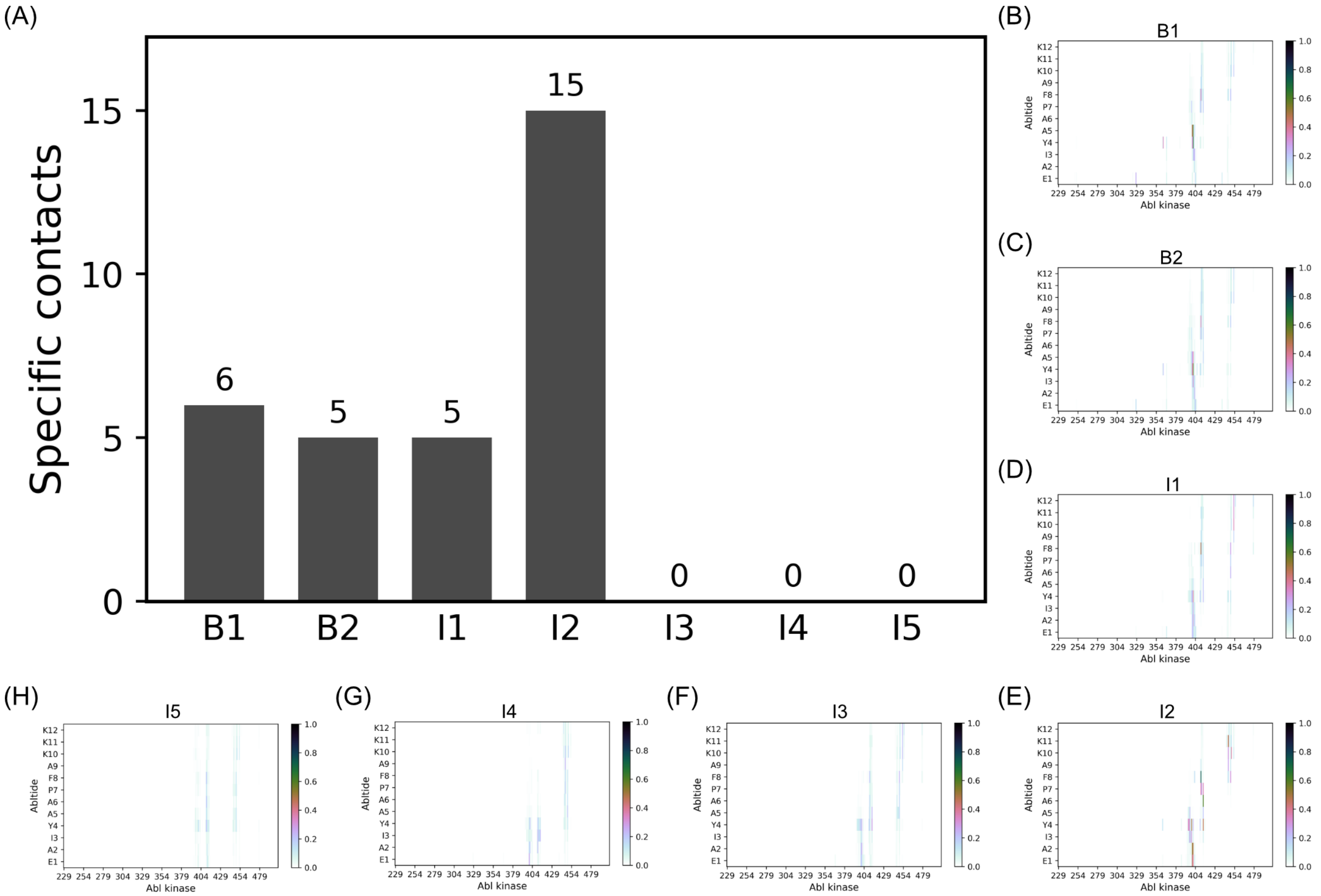


**Fig. S6.** Residue-pair contacts between Abl kinase and Abltide in the states of the bound and intermediate regions. (A) Number of specific contacts (residue pairs with a contact probability above 0.3) in each state. (B–H) Contact probability maps for B1, B2, I1, I2, I3, I4, and I5, respectively.

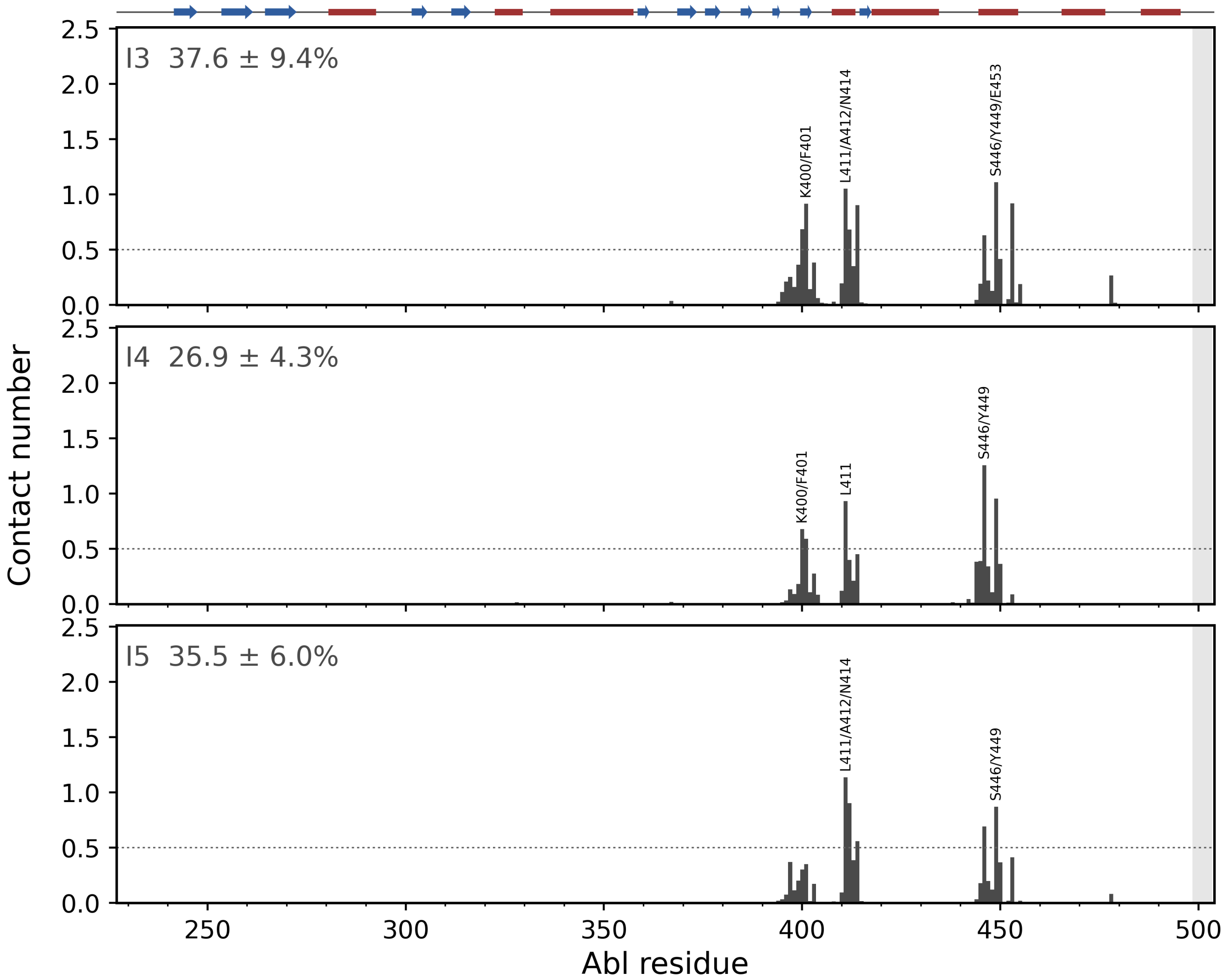


**Fig. S7.** Contact number of each Abl kinase residue with Abltide in each of the three intermediate states I3–I5. Labeling and the shaded region are defined as in Figure S5. The percentage in each panel gives the MBAR-weighted population of the state relative to I3–I5, with its standard error.

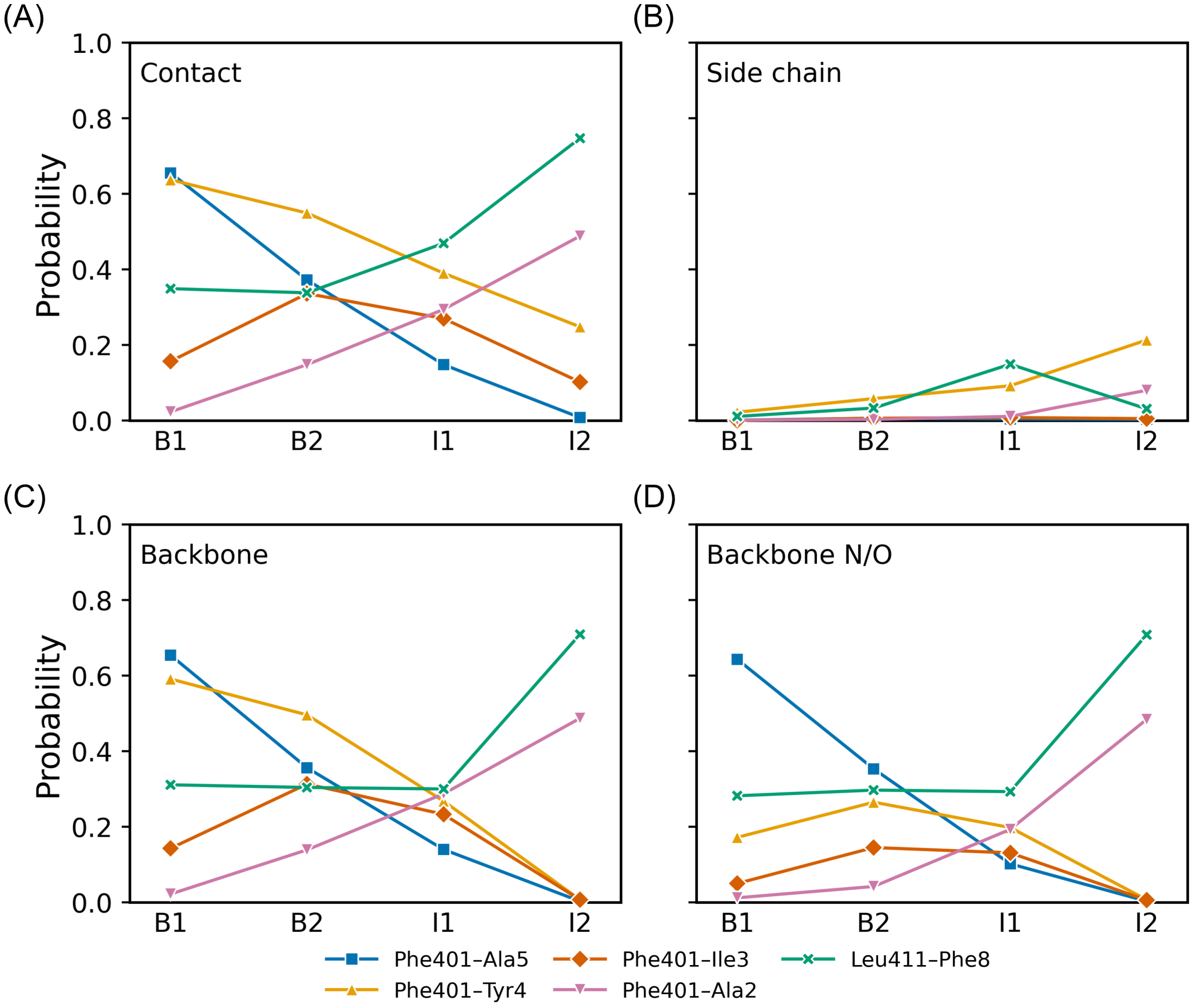


**Fig. S8.** Decomposition of selected Abl-Abltide contacts into side-chain and backbone components in the B1, B2, I1, and I2 states. The five residue pairs shown are Phe401–Ala5, Phe401–Tyr4, Phe401–Ile3, Phe401–Ala2, and Leu411–Phe8. (A) Contact probability of each residue pair, corresponding to the values listed in Table S1. (B–D) Contact probabilities obtained when the distance is evaluated over the side-chain heavy atoms (B), the backbone heavy atoms (C), and the backbone N and O atoms (D) of both residues. Lines connect the states in the order of the progressive register shift (Figure 5 of the main text) and do not represent a temporal sequence.

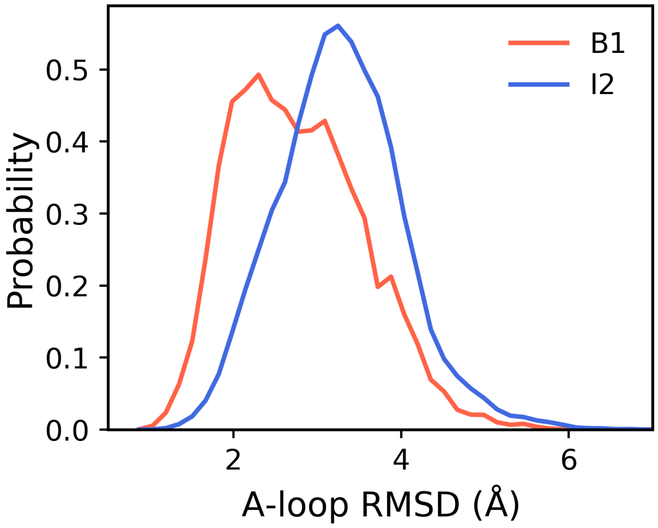


**Fig. S9.** Probability distributions of the A-loop RMSD in the B1 (red line) and I2 (blue line) states. RMSD values were calculated with respect to the initial structure of Abl kinase in the active conformation, prepared from the X-ray crystal structure (PDB ID: 2G2I) (1).

**Table S1. Specific contacts between Abl kinase and Abltide in the B1, B2, I1, and I2 states. Each row lists a residue pair whose contact probability exceeds 0.3 in at least one of the four states, with such values set in bold. The number in parentheses gives the standard error in the last digit (e.g., 0.34(7) denotes 0.34 $\pm$ 0.07) and is omitted where it rounds to zero.**

| Abl | Abltide | B1 | B2 | I1 | I2 |
|---|---|---|---|---|---|
| Asp363 | Tyr4 | **0.34**(7) | 0.21(4) | 0.00 | 0.00 |
| Ala395 | Tyr4 | 0.00 | 0.02 | 0.06(2) | **0.33**(6) |
| His396 | Tyr4 | 0.00 | 0.02 | 0.09(3) | **0.36**(6) |
| Ala399 | Ile3 | 0.01 | 0.02(1) | 0.06(2) | **0.30**(6) |
| Ala399 | Tyr4 | 0.02(1) | 0.13(1) | 0.19(5) | **0.48**(10) |
| Lys400 | Glu1 | 0.06(2) | 0.08(2) | 0.22(4) | **0.43**(7) |
| Lys400 | Ala2 | 0.01 | 0.08(3) | 0.19(5) | **0.52**(10) |
| Lys400 | Tyr4 | **0.31**(1) | **0.36**(4) | 0.22(3) | 0.07(1) |
| Lys400 | Ala5 | **0.48**(5) | 0.27(5) | 0.08(3) | 0.00 |
| Phe401 | Glu1 | 0.01 | 0.05(1) | 0.20(6) | **0.40**(10) |
| Phe401 | Ala2 | 0.02(1) | 0.15(3) | 0.29(8) | **0.49**(12) |
| Phe401 | Ile3 | 0.16(4) | **0.34**(4) | 0.27(3) | 0.10(3) |
| Phe401 | Tyr4 | **0.64**(3) | **0.55**(3) | **0.39**(5) | 0.25(6) |
| Phe401 | Ala5 | **0.66**(6) | **0.37**(6) | 0.15(5) | 0.01 |
| Leu411 | Pro7 | 0.22(3) | 0.22(4) | 0.17(4) | **0.37**(5) |
| Leu411 | Phe8 | **0.35**(3) | **0.34**(4) | **0.47**(6) | **0.75**(9) |
| Asn414 | Tyr4 | 0.05(3) | 0.09(1) | 0.22(6) | **0.47**(8) |
| Asn414 | Ala6 | 0.06(5) | 0.12(1) | 0.12(4) | **0.61**(11) |
| Asn414 | Pro7 | 0.18(2) | 0.16(2) | 0.15(2) | **0.35**(7) |
| Ser446 | Lys11 | 0.05(1) | 0.02(1) | 0.01 | **0.47**(12) |
| Glu450 | Lys10 | 0.14(1) | 0.14(1) | 0.15(3) | **0.38**(8) |
| Glu453 | Lys10 | 0.19(4) | 0.17(3) | **0.35**(4) | 0.12(2) |
| Glu453 | Lys11 | 0.14(1) | 0.13(2) | **0.32**(3) | 0.07(2) |
| Glu453 | Lys12 | 0.13(1) | 0.16(3) | **0.30**(6) | 0.06(2) |
